# A CRISPR-based yeast two-hybrid system for investigating RNA-protein interactions

**DOI:** 10.1101/139600

**Authors:** Evan P. Hass, David C. Zappulla

## Abstract

Despite the great importance of RNA-protein interactions in cells, there is a very limited set of approaches available for identifying proteins that bind to a specific RNA. We report here combining the use of CRISPR technology with the yeast two-hybrid protein-protein interaction system in order to create an advantageous method for investigating RNA-protein interactions. In this <u>C</u>RISPR-<u>a</u>ssisted <u>R</u>NA/<u>R</u>BP <u>y</u>east (CARRY) two-hybrid system, an RNA of interest is targeted to the promoters of standard yeast two-hybrid reporter genes by fusing it to the CRISPR guide RNA in a strain expressing catalytically deactivated Cas9 (dCas9). If the promoter-tethered RNA binds to a protein fused to Gal4 transcriptional activation domain (GAD), then the reporter genes become transcribed, just as in the standard protein-protein yeast two-hybrid assay. We used the CARRY two-hybrid system to analyze MS2 bacteriophage RNA hairpin binding to the MS2 coat protein (MCP). We tested MS2 hairpin mutants with a range of biochemically determined binding affinities for MCP and found that CARRY two-hybrid detected all binding interactions with dissociation constants ≤300 nM. In summary, this new CRISPR-based yeast two-hybrid system provides an easily operable, much-needed new tool for identifying proteins that bind to a particular RNA.

## INTRODUCTION

RNA-binding proteins are integral to the function of RNAs. Many RNA functions are mediated by associated proteins (e.g., chromatin modification by lncRNA-bound enzymes, recruitment of telomerase RNA to telomeres by protein subunits of telomerase). As for functional RNAs that ultimately act protein-independently (e.g., peptide-bond formation by ribosomal RNA, mRNA splicing by spliceosomal RNA), these transcripts still require associated proteins for their proper folding, processing, modification, stabilization, and localization. Because so many cellular RNA-protein interactions remain unknown, it is advantageous to pursue their discovery using high-throughput approaches. The advent and continual improvement of high-throughput DNA sequencing technology has led to the development of many powerful techniques, such as RIP-seq and CLIP-seq, which can be used to identify the full repertoire of RNAs bound by a protein of interest. However, relatively fewer protocols exist for identifying proteins that bind to a particular RNA. Most available techniques involve RNA pull-down followed by protein identification via mass spectrometry, which requires large amounts of starting material, meaning that low-abundance RNA-protein complexes are difficult to study with this approach^1^.

In an effort to address the relative dearth of techniques for identifying binding partners for a given RNA, we have developed a novel technique: <u>C</u>RISPR-<u>a</u>ssisted <u>R</u>NA/<u>R</u>BP <u>y</u>east (CARRY) two-hybrid (Figure 1A). Like the original yeast two-hybrid assay^2,3^, CARRY two-hybrid interrogates binding between two biological macromolecules by tethering one to the promoter of a reporter gene and fusing the second to a transcriptional activation domain. Expression of the reporter gene occurs when there is binding between the two macromolecules. Unlike the original yeast two-hybrid system, instead of tethering a protein of interest to the promoter by fusing it to a DNA-binding domain, CARRY two-hybrid tethers an RNA of interest. RNA tethering is achieved using the *Streptococcus pyogenes* CRISPR machinery. While the CRISPR/Cas9 system has commonly been co-opted for the purpose of making targeted cuts in DNA, nuclease-deactivated Cas9 (dCas9) can target an RNA or protein of interest to a specific genomic locus by fusing it to the CRISPR single-guide RNA (sgRNA) or to Cas9, respectively^4,5^. CARRY two-hybrid uses the former of these two strategies to target an RNA of interest to a shared sequence at the promoters of the yeast two-hybrid reporter genes, *HIS3* and *LacZ*. These reporter genes are then activated if a protein that has been fused to the Gal4 transcriptional activation domain (GAD) binds to the promoter-tethered RNA (Figure 1A).

**Figure 1.**
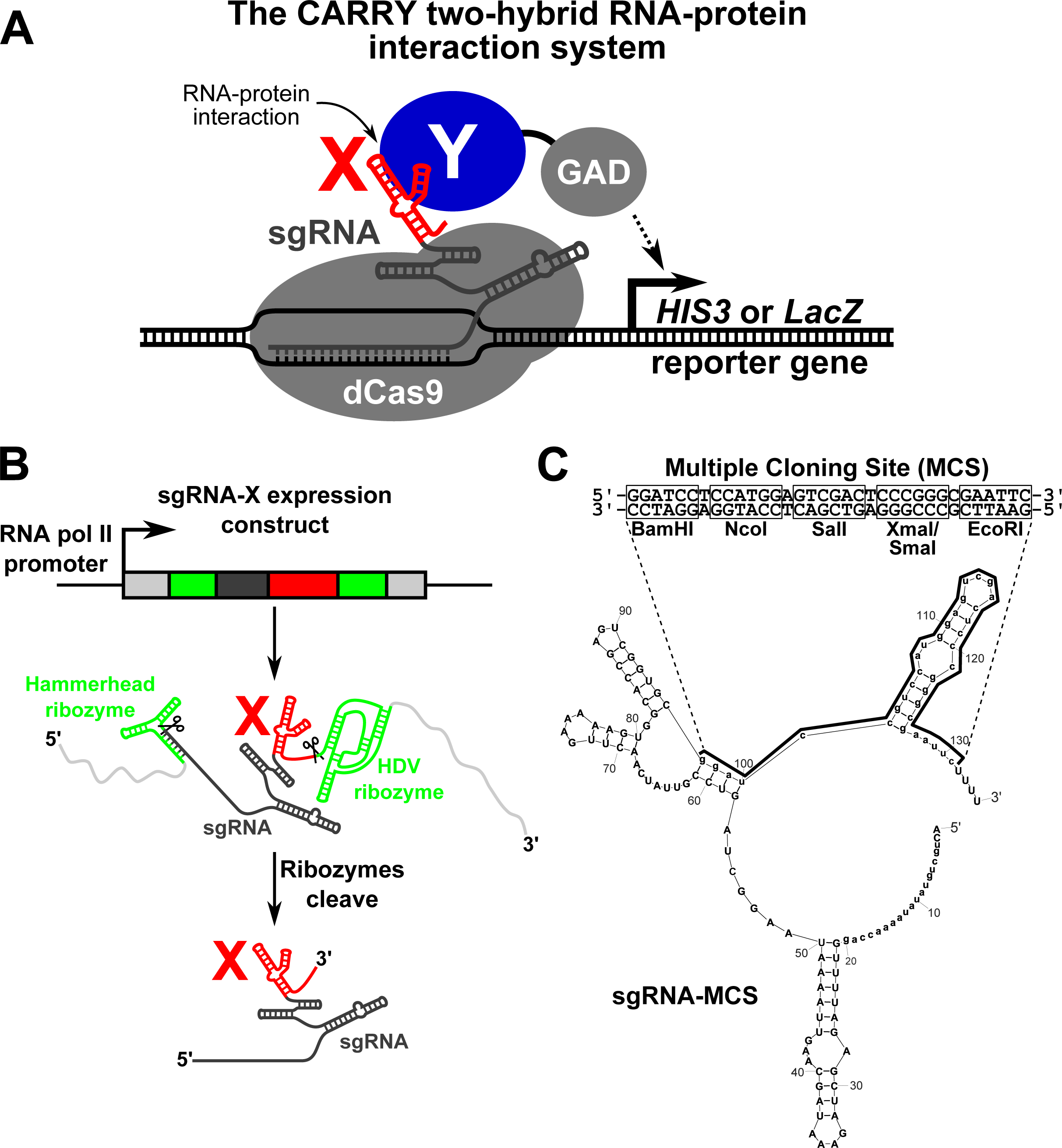
The CARRY two-hybrid assay for interrogating RNA-protein interactions. (**A**) Schematic of the CARRY two-hybrid system. An RNA of interest (“X”, red) is fused to a CRISPR single guide RNA (sgRNA), which is targeted to the promoters of the reporter genes *HIS3* and *LacZ* with assistance from the nuclease-deactivated *Streptococcus pyogenes* Cas9 protein (dCas9). If the RNA of interest fused to the sgRNA binds to the protein (“Y”, blue) fused to the Gal4 activation domain (GAD), the transcription of the reporter gene is activated. (**B**) A schematic of the “RGR” sgRNA expression cassette (adapted from ref. 4 and originally developed by ref. 12). The hybrid sgRNA is expressed from an RNA polymerase II promoter, flanked by the hammerhead and HDV ribozymes (green). Once transcribed, the ribozymes catalyze self-cleavage of the RNA, processing the mature hybrid sgRNA out of the longer transcript. (**C**) The hybrid sgRNA plasmid used in CARRY two-hybrid contains a multiple-cloning site (MCS) that forms a hairpin when transcribed into RNA. Shown is an *Mfold* prediction of the secondary structure of the sgRNA-MCS, in which the guide of the sgRNA was forced to be single-stranded. The MCS RNA sequence is bracketed, and its DNA sequence is shown underneath, with its five unique restriction sites annotated.

Here, we show that CARRY two-hybrid works. The yeast two-hybrid reporter genes are activated contingent on binding between an sgRNA-fused RNA and GAD-fused protein. Furthermore, CARRY two-hybrid is specific, and our results show that it is sufficiently sensitive to detect RNA-protein interactions with near-micromolar dissociation constants. We expect that CARRY two-hybrid will prove to be a very useful new tool for the identification and characterization of RNA-protein interactions.

## RESULTS

### Design and construction of a yeast two-hybrid system to study RNA-protein binding

We constructed the yeast strain used for CARRY two-hybrid, “CARRYeast (1.0),” by integrating a dCas9 expression cassette^4^ in the genome of a previously published yeast two-hybrid strain, L40, which contains the reporter genes *HIS3* and *LacZ* with 4 and 8 LexA-binding sites inserted in their promoters, respectively^6^. While several adaptations of the CRISPR/Cas9 system used in *S. cerevisiae* express the sgRNA from an RNA polymerase III promoter^7,8^, we chose to express the hybrid sgRNA for CARRY two-hybrid using an RNA polymerase II promoter, since RNA polymerase III transcription can be terminated by even a relatively short poly(U) tract^9^, whereas RNA polymerase II termination signals are more complex and therefore should be rarer^10,11^. Thus, because premature termination of transcription in the middle of the hybrid sgRNA would make the CARRY two-hybrid system unusable, RNA polymerase II ultimately imposes fewer restrictions than RNA polymerase III on the RNA sequences that can be tested in this system.

In order to express the hybrid sgRNA, we modified a previously published RNA polymerase II sgRNA expression construct^4^ (Figure 1B). Because the mRNA promoter and terminator introduce extraneous sequence at the 5′ and 3′ ends of the expressed RNA (e.g., the poly(A) tail), we chose to use a construct that employs a ribozyme-guide RNA-ribozyme (RGR) cassette for sgRNA processing^12^. In an RGR cassette, an sgRNA is flanked by the hammerhead and HDV ribozymes that self-cleave, thus excising the sgRNA from the longer initial transcript *in vivo*. We cloned this RNA polymerase II RGR sgRNA expression cassette into a centromeric yeast vector and changed the guide sequence at the 5¢ end of the sgRNA to target the RNA to the LexA-binding sites upstream of both the *HIS3* and *LacZ* reporter genes. Finally, in order to facilitate the cloning of diverse RNA domains into this hybrid sgRNA expression vector, we inserted a multiple-cloning site (MCS) containing five unique, commonly used restriction-enzyme cleavage sites near the 3′ end of the sgRNA, four nucleotides 5′ of the HDV ribozyme cleavage site. Because at least some of the MCS will ultimately be part of the transcribed hybrid sgRNA (depending on the restriction sites used for subcloning), it was designed to form a hairpin when transcribed, making it less likely to pair with and disrupt folding of the inserted RNA of interest. In an *Mfold* computational prediction of the sgRNA-MCS secondary structure, in which the guide sequence was forced to be single-stranded, most of the MCS sequence is indeed predicted to form a hairpin, as designed (Figure 1C). Although the first four nucleotides of the MCS sequence are predicted to pair with part of the sgRNA rather than with the last four nucleotides of the MCS sequence, these few predicted base pairs (one of which is a G•U pair) apparently did not prevent the expected tethering of the sgRNA to its target sites by dCas9 based on reporter-gene activation results (see below).

### CARRY two-hybrid can specifically detect the MS2-MCP interaction

We first sought to test the CARRY two-hybrid system with a well-understood RNA-protein interaction, such as the MS2 bacteriophage’s RNA hairpin binding to the phage’s coat protein. We cloned the MS2 RNA hairpin mutant, U-5C – which binds the MS2 coat protein more tightly than the wild-type hairpin^13^ – into the sgRNA expression vector, and we also cloned a tandem dimer of the MS2 coat protein (MCP_2_) into a standard vector for expression of Gal4-activating domain (GAD) fusion proteins in the yeast two-hybrid system, pGAD424^3,14^. These plasmids were then transformed into CARRYeast (1.0), and expression of *HIS3* and *LacZ* were assessed by growth of cells on medium lacking histidine and by a colorimetric assay, respectively. When both the sgRNA-U-5C MS2 hybrid RNA and the GAD-MCP_2_ hybrid protein were expressed, expression of both *HIS3* and *LacZ* was strongly induced (Figure 2A, third row; Figure 2B, bottom right). Importantly, activation was dependent on the MS2 hairpin being fused to the sgRNA (Figure 2A, rows 1 and 2; Figure 2B, top panels), and MCP_2_ being fused to GAD (Figure 2A, rows 2 and 4; Figure 2B, left panels). This indicates that the CARRY two-hybrid system is able to detect RNA-protein interactions, and that it does so specifically. Furthermore, while we observed some low-level background expression of the *LacZ* reporter gene in some negative controls, as is often observed in the standard yeast two-hybrid system using the CARRYeast (1.0)’s parent strain L40^15,16^, the *HIS3* reporter gene (which is the reporter gene to be used for forward-genetic selection of GAD fusion-protein libraries), consistently showed no background signal with negative controls.

**Figure 2.**
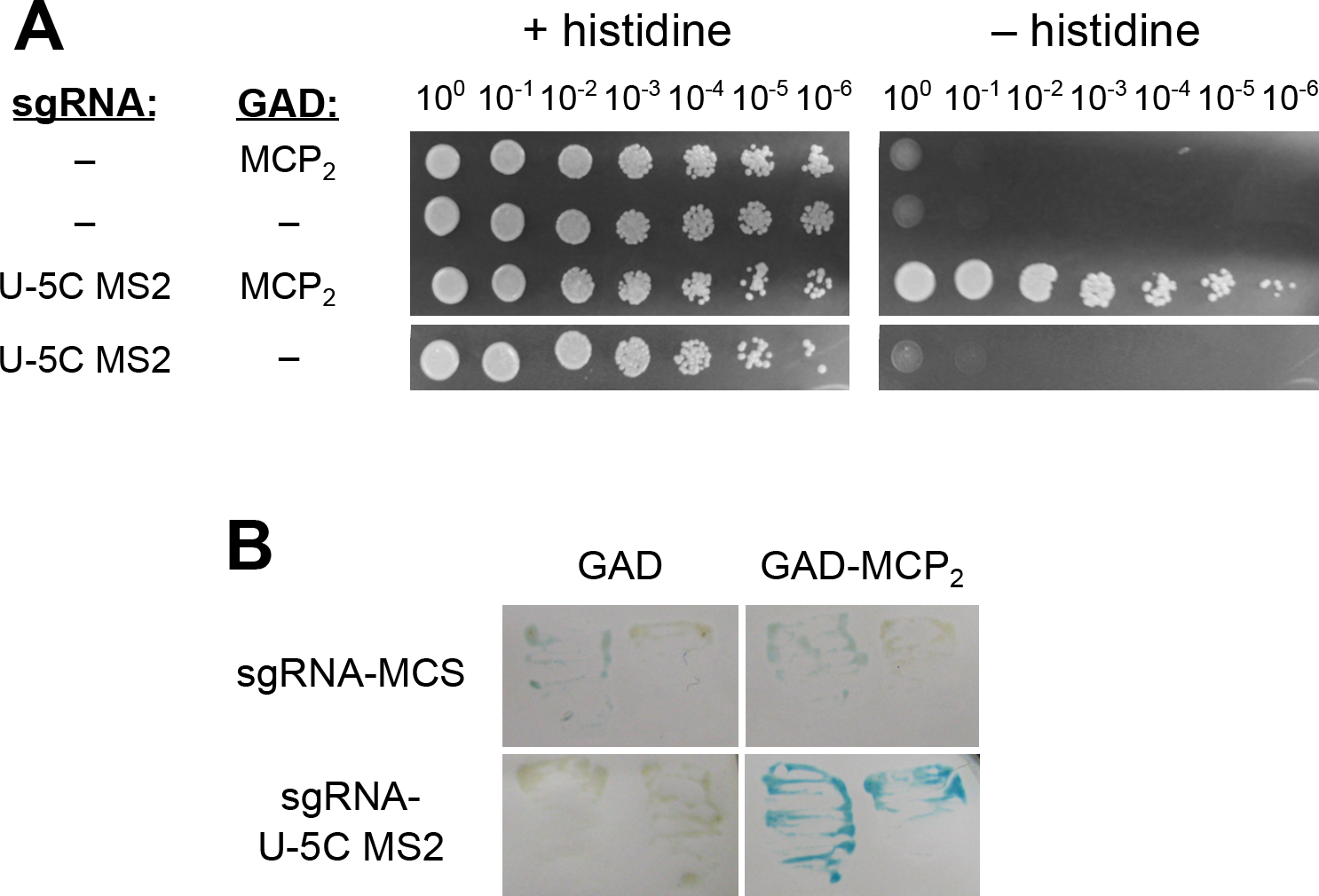
The MS2-MCP interaction strongly activates the *HIS3* and *LacZ* reporter genes of the CARRY two-hybrid system. (**A**) The *HIS3* reporter gene in CARRYeast (1.0) is activated strongly and specifically by the MS2-MCP interaction. Yeast cells were grown to saturation in liquid culture. Cells from the undiluted culture and from six 10-fold serial dilutions (as indicated above the images) were spotted to, and grown on, medium with or without histidine. In the columns on the left, minus signs denote that the sgRNA or GAD were not fused to any RNA or protein, respectively. In the case of the sgRNA, this means that it contained the MCS sequence shown in Figure 1C. (**B**) The *LacZ* reporter gene in CARRYeast (1.0) is activated strongly and specifically by the MS2-MCP interaction. Yeast were grown overnight, lysed on a nitrocellulose filter with liquid nitrogen, and exposed to X-gal, as described in Methods. The filter was left at 30°C overnight for color to develop until the filter had dried out and the reaction had stopped. Pairs of yeast patches are biological replicate samples.

### CARRY two-hybrid can detect RNA-protein interactions with near-micromolar dissociation constants

Next, to test the sensitivity of the CARRY two-hybrid system, we replaced the U-5C MS2 hairpin with the wild-type MS2 hairpin and several biochemically characterized mutants of the MS2 hairpin with a range of weaker binding affinities for the MS2 coat protein^13^. In our tests of these MS2 hairpin mutants, *HIS3* and *LacZ* were activated by interactions several orders of magnitude weaker than the U-5C MS2 interaction with MCP (K_d_ ≈ 20 pM) (Figure 3B, C). For the wild-type MS2 hairpin (K _d_ ≈ 3 nM) and the C-14A/U-12A/A1U/G3U MS2 hairpin (K _d_ ≈ 45 nM, hereafter referred to as the AU-helix MS2 hairpin), activation of the *HIS3* and *LacZ* reporters appeared just as strong as that for the U-5C hairpin. For the A-7C hairpin (K _d_ ≈ 300 nM), activation of the *HIS3* reporter was detectable when undiluted culture was spotted on medium lacking histidine (Figure 3B, row 5), whereas no activation of the *LacZ* reporter was observed (Figure 3C, lower left panel). Finally, the A-7U hairpin, which binds to MCP with a dissociation constant ≥10 μM, did not activate either *HIS3* or *LacZ*.

**Figure 3.**
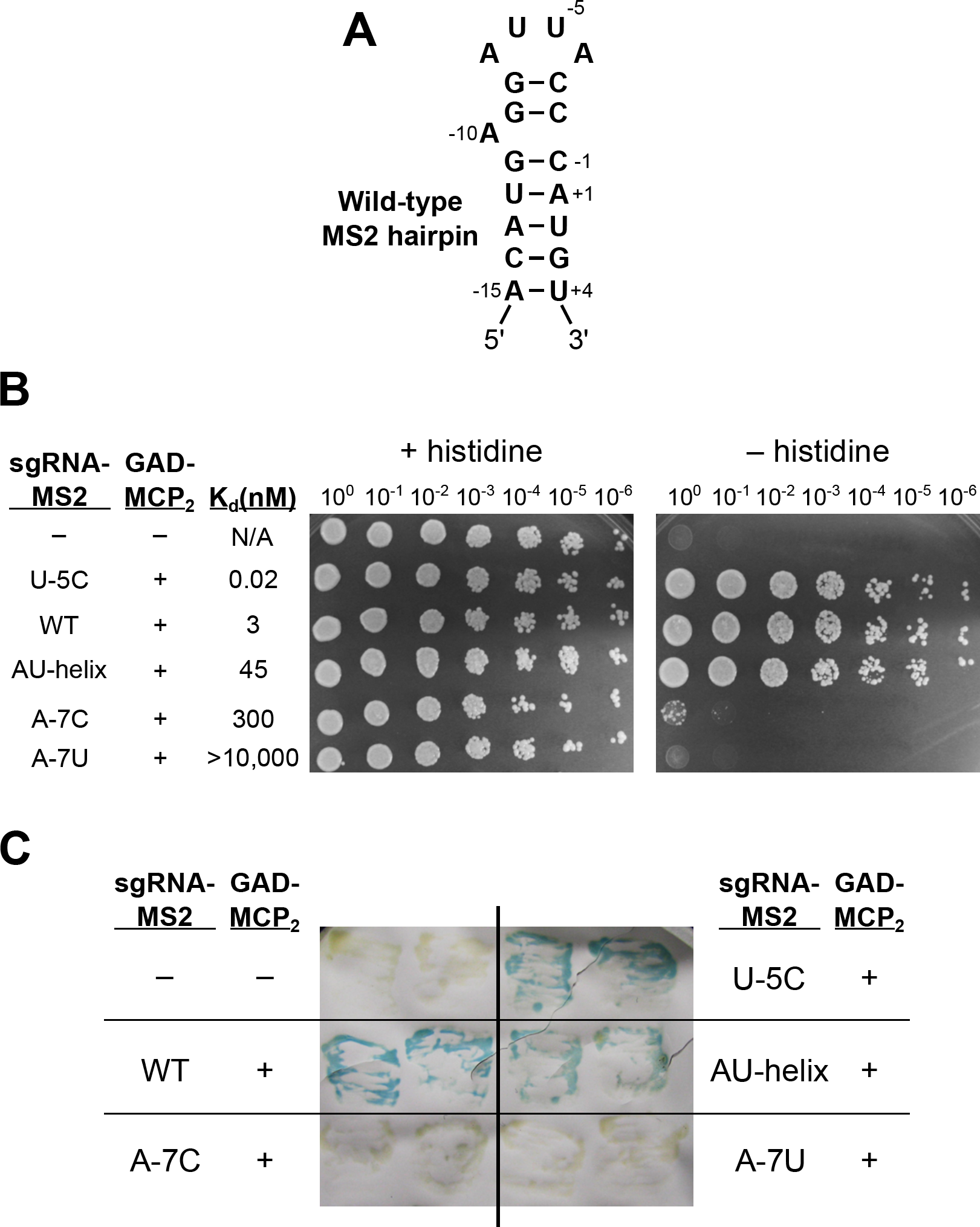
The sensitivity of the CARRY two-hybrid system: detection of RNA-protein binding interactions with near-micromolar dissociation constants. (**A**) The secondary structure of the wild-type MS2 RNA hairpin. Nucleotide numbering is based on the AUG start codon of the MS2 bacteriophage’s replicase gene and serves as the basis for the nomenclature of MS2 point mutants. (**B**) The *HIS3* reporter gene in CARRYeast (1.0) is activated robustly by MS2-MCP interactions with dissociation constants as high as 300 nM. Yeast were grown as in Figure 2A on solid media containing or lacking histidine. The dissociation constants reported here and in the text were calculated using association constants reported previously^13^. “AU-helix” refers to the C-14A/U-12A/A1U/G3U MS2 hairpin quadruple mutant. (**C**) The *LacZ* reporter gene in CARRYeast (1.0) is activated by MS2-MCP interactions with dissociation constants as high as 45 nM. Yeast were grown, lysed, and exposed to X-gal as in Figure 2B. Pairs of yeast patches are biological replicate samples. In Figures 3B and 3C, as in Figure 2A, minus signs denote that no RNA or protein was fused to the sgRNA or GAD, respectively. In this figure, an earlier design of the sgRNA MCS sequence, MCSv0.5, was used as a negative control (see Methods).

Next, to test if we could increase the sensitivity of the CARRY two-hybrid system, we subcloned the sgRNA expression cassette from a ∼single-copy centromeric plasmid to a high-copy 2µ (2-micron) plasmid and re-tested activation for several of the MS2 hairpin mutants. Although expression of the hybrid sgRNA from the high-copy plasmid could not increase the already-maximal *HIS3* activation for the U-5C or AU-helix mutant MS2 hairpins (Figure 4A, compare row 2 with 6 and 3 with 7), in contrast, the activation of the *HIS3* reporter was increased ∼10,000-fold for the A-7C MS2 RNA hairpin (Figure 4A, compare rows 4 and 8) compared to when the sgRNA was expressed from a low-copy plasmid. Importantly, the negative controls — either expressing the sgRNA alone or GAD alone when using the high-copy plasmid — did not result in any detectable *HIS3* activation (Figure 4B) or *LacZ* activation (data not shown). In contrast to results with the *HIS3* reporter gene, activation of the *LacZ* reporter was not visibly increased by expressing the hybrid sgRNA from a high-copy plasmid (Figure 4C). Thus, in summary, although the *LacZ* reporter in the CARRY two-hybrid system is not very responsive, the *HIS3* reporter is sensitive, with low background and substantial dynamic range, making it highly useful as an *in vivo* indicator of RNA-protein binding.

**Figure 4.**
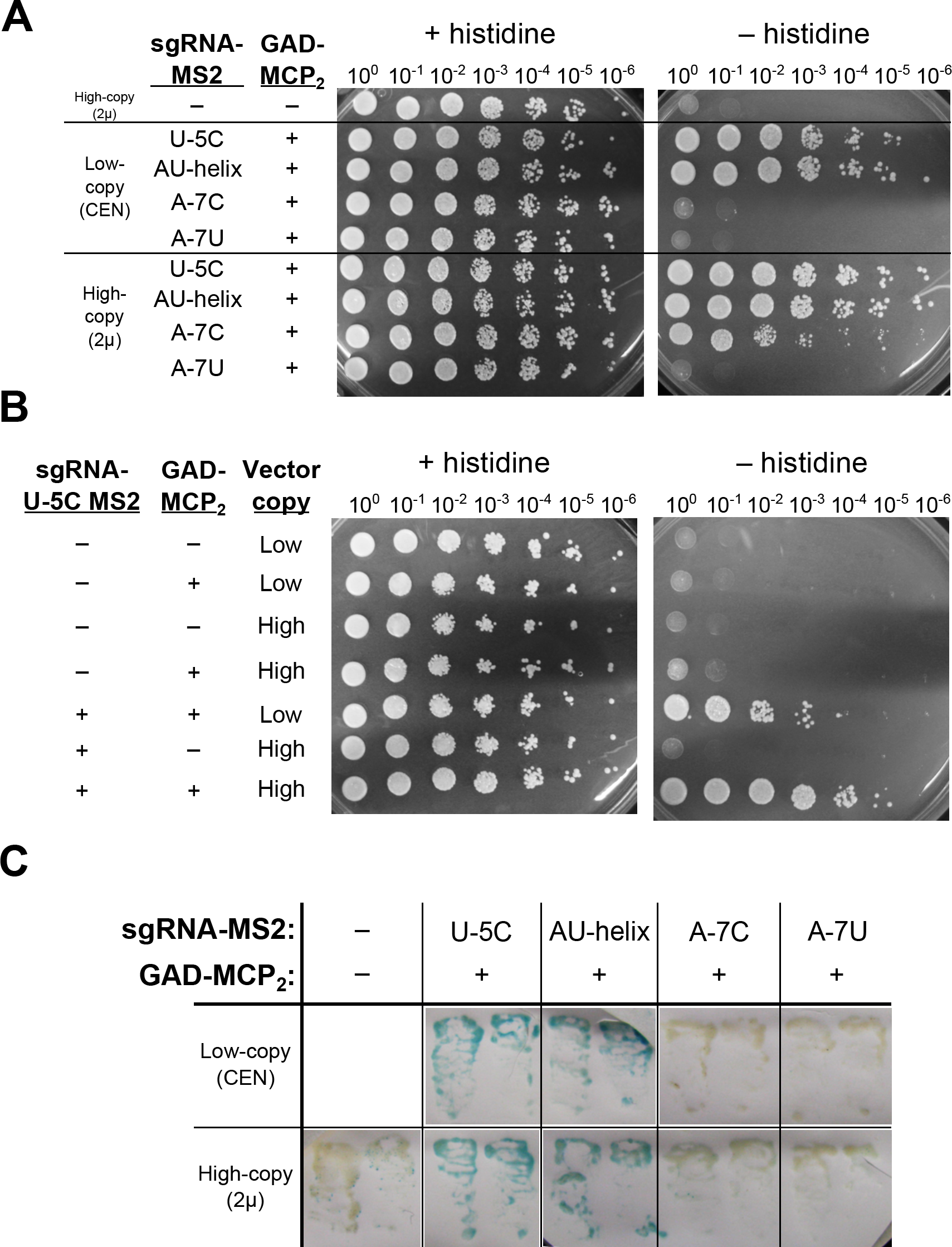
Expression of the hybrid sgRNA from a high-copy plasmid increases assay sensitivity. (**A and B**) Expression of the hybrid sgRNA from a high-copy plasmid increases *HIS3* reporter gene activation, allowing detection of low-affinity RNA-protein interactions. Yeast were grown as in Figure 2A on solid medium with or without histidine. Despite the increase in sensitivity of CARRY two-hybrid permitted by the increase in expression on 2µ plasmids, background signal was not increased. (**C**) Expression of the hybrid sgRNA from a high-copy (2µ) plasmid did not perceptibly increase activation strength of the *LacZ* reporter gene. Yeast were grown, lysed, and exposed to X-gal substrate for LacZ, as in Figure 2B. Pairs of yeast patches shown are biological-replicate samples. In Figure 4, as in the previous figures, a minus signs denote that no RNA or protein was fused to the sgRNA or GAD, respectively; i.e., pCARRY1, pCARRY2, and pGAD424 “vector-only” negative controls.

## DISCUSSION

We have developed a new assay for investigating RNA-protein interactions, “CARRY two-hybrid,” that combines CRISPR/dCas9 RNP-mediated targeting of RNA to a specific DNA sequence with the highly effective yeast two-hybrid protein-protein interaction assay. As evidenced by tests we performed using CARRY two-hybrid to analyze bacteriophage MS2 hairpin binding to MS2 coat protein, this new assay can detect RNA-protein interactions *in vivo* with high specificity (i.e., virtually no background signal for the *HIS3* reporter gene) and can detect interactions with near-micromolar dissociation constants.

Given the simplicity of the CARRY two-hybrid system and the ease with which it has functioned in our hands thus far, we expect that it will prove to be a highly effective method for dissecting known RNA-protein interfaces, as well as for the discovery of new RNA-protein interactions. We have constructed a vector with a multiple-cloning site to facilitate fusing an RNA of interest to the sgRNA (see Figure 1C). The RNA polymerase II promoter allows CARRY two-hybrid to be used to study a large variety of RNA-encoding DNA sequences, and the self-cleaving ribozymes in the initial transcript RNA “bait” in the two-hybrid system trim extraneous sequences from the 5¢ and 3¢ ends^4,12^ (Figure 1B). Additionally, because the CARRY two-hybrid assay is built upon the well-established protein-protein yeast two-hybrid system^17^, the existing GAD fusion libraries constructed by labs and companies can now also be used for studying proteins binding to RNA.

CARRY two-hybrid is similar to the yeast “three-hybrid” system in the sense that the three-hybrid method also assays for RNA-protein interactions by building upon the basic principles underlying the original yeast two-hybrid assay. The three-hybrid system, published over 15 years ago, employs the well-characterized, high-affinity MS2-MCP RNA-protein interaction (or RRE-RevM10 from HIV^18^) to tether RNAs of interest to reporter-gene promoters by way of fusing them to the MS2 hairpin, while also appending MCP to a specific DNA-binding protein domain; thus, there is a total of three hybrid molecules^19^. However, there has been limited success using the three-hybrid system, as evidenced by the relative paucity of publications referencing use of three-hybrid. Although we have yet to directly compare the capabilities of CARRY two-hybrid with those of yeast three-hybrid, we anticipate that CARRY two-hybrid is likely to prove to be even more useful. The recruitment of the Gal4 activating domain to the reporter genes in yeast three-hybrid necessitates three different binding interactions (e.g., DNA LexA sites • LexA_DBD_–MCP • MS2 RNA–X • Y–GAD). In contrast, the CARRY two-hybrid system uses CRISPR/dCas9 to directly target RNA to DNA. By reducing the number of stable binding events required for activating reporter genes to two, as well as other features that promote efficiency and robustness described above, CARRY two-hybrid is likely to be more effective at detecting RNA-protein interactions. Future studies will show if this bears out.

## METHODS

### Construction of yeast strains and plasmids for CARRY two-hybrid

CARRYeast (1.0) was generated by modifying the yeast two-hybrid strain L40 (*MATa his3*Δ*200 trp1-901 leu2-3,112 ade2 LYS2::(4LexAop-HIS3) URA3::(8LexAop-LacZ)*)^6^. First, yeast cells were transformed with linearized pJZC518^4^ containing a cassette for expression of the *S. pyogenes* dCas9 in *S. cerevisiae*, *C. glabrata LEU2* selectable marker, and homology arms for integration at the *S. cerevisiae LEU2* locus. In the resulting yeast strain, the *C. glabrata LEU2* selectable marker was subsequently deleted using a cassette generated using pFA6a-KanMX6^20^.

The sgRNA expression vectors, pCARRY1 and pCARRY2, were based on pJZC625^4^. This plasmid contains a ribozyme-guide RNA-ribozyme (RGR) cassette^12^. The sgRNA in pJZC625 contained a guide sequence targeted to the TET operator and a U-5C MS2 hairpin inserted 4 nucleotides before the HDV ribozyme cut site. The RGR cassette is flanked by the *S. cerevisiae ADH1* promoter and the *C. albicans ADH1* terminator. To generate pCARRY1, pJZC625 was digested with ApaI and BglII, and the full expression cassette was cloned into pRS414^21^ that had been digested with ApaI and BamHI. Second, the guide sequence of the sgRNA was changed to target the LexA operator sequence ACTGCTGTATATAAAACCAG, which is followed by a protospacer adjacent motif (PAM) with sequence TGG in the LexA operators present in CARRYeast (1.0). Additionally, in order to maintain base-pairing in the H1 stem of the hammerhead ribozyme of the RGR cassette (the 3′ half of which consists of the first 6 nucleotides of the sgRNA guide sequence), the sequence of the 5′ half of the H1 stem was changed to AGCAGT. Third, the MS2 hairpin was replaced with GGATCCCATGGGTCGACCCCGGGAATTC, an earlier-designed version of the hairpin-forming multiple-cloning site sequence (MCSv0.5). This sequence was later replaced with the MCS sequence shown in Figure 1C (GGATCCGTCCATGGAGTCGACTCCCGGGCGAATTC), generating pCARRY1. This modified version of the original RGR expression construct was then subcloned into pRS424^22^ using KpnI and SpeI to generate pCARRY2. The original U-5C MS2 hairpin sequence present in pJZC625 (GCGCACATGAGGATCACCCATGTGC) and mutants thereof were cloned into pCARRY1 and pCARRY2 using BamHI and EcoRI.

The vector used to express the GAD-MCP_2_ fusion protein, pDZ982, was cloned using pGAD424^14^. DNA encoding a tandem MCP dimer and an N-terminal linker (i.e., ultimately between GAD and MCP_2_ in the final plasmid) with amino-acid sequence GGGR was PCR amplified from the plasmid pDZ349^23^ and cloned into pGAD424 using XmaI and PstI. Both MCP monomers contain the N55K mutation, reported to strengthen binding to the MS2 hairpin ∼10-fold^24^, while the first monomer also contains the incidental mutations K57R and I104V.

### *HIS3* reporter gene spot assay

Expression of the *HIS3* reporter gene in CARRYeast (1.0) was assayed by first growing yeast in liquid culture (using minimal medium lacking tryptophan and leucine) to saturation overnight. 100-μL aliquots were taken from these cultures and used to make six 10-fold serial dilutions of the culture. 5 μL of the undiluted aliquot and each serial dilution were pipetted onto both solid –Trp –Leu and –Trp –Leu –His minimal media. These spotted cells were then incubated for two days at 30°C and photographed.

### *LacZ* reporter gene assay

Colorimetric *LacZ* reporter gene expression assays were performed as described previously^14^. Briefly, expression of the *LacZ* reporter gene in CARRYeast (1.0) was assayed by first streaking the cells as patches on –Trp –Leu medium and incubating for ∼15–24 hours at 30°C. Yeast were then removed from the agar plate by applying and removing a circular nitrocellulose membrane. Yeast attached to the nitrocellulose were lysed by brief submergence in liquid nitrogen. Then, in a petri dish, a piece of Whatman filter paper was wetted with 1.8 mL of 100 mM sodium phosphate buffer pH 7.0 with 10 mM KCl, 1 mM MgSO_4_, and 333 μg/mL X-gal. The nitrocellulose filter was soaked in the X-gal solution by laying it, face up, on top of the Whatman paper, and the petri dish was incubated at 30°C. The color of the lysed yeast cells was monitored and photographed at time intervals over ∼24 hours, or until the dish had dried out and the reaction had stopped.

## ACKNOWLEDGEMENTS

We thank Jesse Zalatan (U. Washington) for reagents. Research reported in this publication was supported by the National Institute of General Medical Sciences of the National Institutes of Health under award number R01GM118757, and startup funds from Johns Hopkins University to DCZ. The content is solely the responsibility of the authors and does not necessarily represent the official views of the National Institutes of Health.

## REFERENCES

1. McHugh, C.A., Russell, P. & Guttman, M. Methods for comprehensive experimental identification of RNA-protein interactions. Genome Biol 15, 203 (2014).

2. Fields, S. & Song, O. A novel genetic system to detect protein-protein interactions. Nature 340, 245–6 (1989).

3. Chien, C.T., Bartel, P.L., Sternglanz, R. & Fields, S. The two-hybrid system: a method to identify and clone genes for proteins that interact with a protein of interest. Proc Natl Acad Sci U S A 88, 9578–82 (1991).

4. Zalatan, J.G. et al. Engineering complex synthetic transcriptional programs with CRISPR RNA scaffolds. Cell 160, 339–50 (2015).

5. Shechner, D.M., Hacisuleyman, E., Younger, S.T. & Rinn, J.L. Multiplexable, locus-specific targeting of long RNAs with CRISPR-Display. Nat Methods 12, 664–70 (2015).

6. Hollenberg, S.M., Sternglanz, R., Cheng, P.F. & Weintraub, H. Identification of a new family of tissue-specific basic helix-loop-helix proteins with a two-hybrid system. Mol Cell Biol 15, 3813–22 (1995).

7. DiCarlo, J.E. et al. Genome engineering in Saccharomyces cerevisiae using CRISPR-Cas systems. Nucleic Acids Res 41, 4336–43 (2013).

8. Laughery, M.F. et al. New vectors for simple and streamlined CRISPR-Cas9 genome editing in Saccharomyces cerevisiae. Yeast 32, 711–20 (2015).

9. Allison, D.S. & Hall, B.D. Effects of alterations in the 3’ flanking sequence on in vivo and in vitro expression of the yeast SUP4-o tRNATyr gene. EMBO J 4, 2657–64 (1985).

10. Tian, B. & Graber, J.H. Signals for pre-mRNA cleavage and polyadenylation. Wiley Interdiscip Rev RNA 3, 385–96 (2012).

11. Porrua, O. & Libri, D. Transcription termination and the control of the transcriptome: why, where and how to stop. Nat Rev Mol Cell Biol 16, 190–202 (2015).

12. Gao, Y. & Zhao, Y. Self-processing of ribozyme-flanked RNAs into guide RNAs in vitro and in vivo for CRISPR-mediated genome editing. J Integr Plant Biol 56, 343–9 (2014).

13. Romaniuk, P.J., Lowary, P., Wu, H.N., Stormo, G. & Uhlenbeck, O.C. RNA binding site of R17 coat protein. Biochemistry 26, 1563–8 (1987).

14. Bartel, P.L., Chien, C.-T., Sternglanz, R. & Fields, S. Using the two-hybrid system to detect protein-protein interactions. in *Cellular Interactions in Development: A Practical Approach* (ed. Hartley, D.A.) 153–179 (Oxford University Press, Oxford, 1993).

15. Zappulla, D.C., Maharaj, A.S., Connelly, J.J., Jockusch, R.A. & Sternglanz, R. Rtt107/Esc4 binds silent chromatin and DNA repair proteins using different BRCT motifs. BMC Mol Biol 7, 40 (2006).

16. Andrulis, E.D. et al. Esc1, a nuclear periphery protein required for Sir4-based plasmid anchoring and partitioning. Mol Cell Biol 22, 8292–301 (2002).

17. Vidal, M. & Fields, S. The yeast two-hybrid assay: still finding connections after 25 years. Nat Methods 11, 1203–6 (2014).

18. Putz, U., Skehel, P. & Kuhl, D. A tri-hybrid system for the analysis and detection of RNA‐‐protein interactions. Nucleic Acids Res 24, 4838–40 (1996).

19. SenGupta, D.J. et al. A three-hybrid system to detect RNA-protein interactions *in vivo*. Proc Natl Acad Sci U S A 93, 8496–501 (1996).

20. Longtine, M.S. et al. Additional modules for versatile and economical PCR-based gene deletion and modification in *Saccharomyces cerevisiae*. Yeast 14, 953–61. (1998).

21. Sikorski, R.S. & Hieter, P. A system of shuttle vectors and yeast host strains designed for efficient manipulation of DNA in *Saccharomyces cerevisiae*. Genetics 122, 19–27. (1989).

22. Christianson, T.W., Sikorski, R.S., Dante, M., Shero, J.H. & Hieter, P. Multifunctional yeast high-copy-number shuttle vectors. Gene 110, 119–22 (1992).

23. Lebo, K.J., Niederer, R.O. & Zappulla, D.C. A second essential function of the Est1-binding arm of yeast telomerase RNA. RNA (2015).

24. Lim, F., Spingola, M. & Peabody, D.S. Altering the RNA binding specificity of a translational repressor. J Biol Chem 269, 9006–10 (1994).

